# Microbial regulation of enteric eosinophils and its impact on tissue remodeling and Th2 immunity

**DOI:** 10.1101/527473

**Authors:** Rodrigo Jiménez-Saiz, Varun C. Anipindi, Heather Galipeau, Yosef Ellenbogen, Roopali Chaudhary, Joshua F. Koenig, Melissa E. Gordon, Tina D. Walker, Talveer S. Mandur, Soumeya Abed, Alison Humbles, Derek K. Chu, Jonas Erjefält, Kjetil Ask, Elena F. Verdú, Manel Jordana

## Abstract

Eosinophils have emerged as multifaceted cells that contribute to tissue homeostasis. However, the impact of the microbiota on their frequency and function at mucosal sites remains unclear. Here, we investigated the role of the microbiota in the regulation of enteric eosinophils. We found that small intestinal (SI) eosinophilia was significantly greater in germ-free (GF) mice compared to specific pathogen free (SPF) controls. This was associated with changes in the production of enteric signals that regulate eosinophil attraction and survival, and was fully reversed by complex colonization. Additionally, SI eosinophils of GF mice exhibited more cytoplasmic protrusions and less granule content than SPF controls. Lastly, we generated a novel strain of eosinophil-deficient GF mice. These mice displayed intestinal fibrosis and were less prone to allergic sensitization as compared to GF controls. Overall, our study demonstrates that commensal microbes regulate intestinal eosinophil frequency and function, which impacts tissue repair and allergic sensitization to food antigens. These data support a critical interplay between the commensal microbiota and intestinal eosinophils in shaping homeostatic, innate and adaptive immune processes in health and disease.

## INTRODUCTION

Eosinophils have traditionally been described as effector inflammatory cells that are protective against parasitic infections but detrimental in allergic disease ^1,2^. New evidence has considerably broadened this paradigm as eosinophils have now been shown to play complex roles in mucosal immunity and tissue remodeling ^3-8^. For example, intestinal eosinophils and eosinophil peroxidase (EPO) are critical for the initiation of Th2 responses (IgE) to food allergens in the gastrointestinal (GI) tract ^6,9^. Furthermore, intestinal accumulation of eosinophils has been associated with the severity of inflammatory bowel disease (IBD)^10,11^. Conversely, intestinal eosinophils secrete large quantities of IL-1 receptor antagonist, thus reducing inflammation in IBD^12,13^. Eosinophils have also been proposed to play a role in homeostatic tissue remodeling as their presence is increased at tissues with a high turnover such as the intestine and the uterus ^4,14^. Eosinophils are equipped with damage-sensing receptors the ligation of which leads to the production of factors involved in tissue repair (*eg.* TGF-α and -β, vascular-endothelial growth factor (VEGF), *etc.*) ^15^. Hence, deciphering the factors that regulate eosinophils in the GI mucosa is relevant to understanding their role in health and disease.

There is growing evidence on the role of the microbiota in regulating immune responses and maintaining intestinal homeostasis ^16,17^. Studies in germ-free (GF) mice have demonstrated that the microbiota is crucial for the maturation of the adaptive immune system in the small intestine (SI) ^18^. For example, GF mice have fewer CD4^+^ T cells, intraepithelial lymphocytes, and lower IgA-producing plasma cells in the lamina propria (LP) of the SI as compared to specific pathogen free (SPF) mice ^16,17,19,20^. Yet, knowledge on the effect of the microbiota on innate cells that are indigenous to the intestinal tract, such is eosinophils, is scarce ^13,21^.

Here we examined the impact of the microbiota on eosinophil frequency and function in the SI. We found that eosinophil frequency was enriched along the SI of GF mice relative to SPF controls. This was associated with local changes in the production of signals involved in the attraction, retention and survival of eosinophils. This relative eosinophilia was also observed in other mucosal sites, but not in sterile tissues, and was corrected by repletion of microbiota through co-habitation with altered Schaedler flora (ASF)- or SPF-mice. Additionally, SI eosinophils of GF mice exhibited more cytoplasmic protrusions and less granule content than SPF controls; this was consistent with the lower EPO levels detected in intestinal homogenates from GF mice. Lastly, we generated a novel strain of eosinophil-deficient (ΔdblGATA1) mice on a GF background. In this system, the absence of eosinophils was associated with increased collagen accumulation in the submucosa as well as reduced allergic sensitization. This study illustrates a novel role for the microbiota in regulating mucosal eosinophils and tissue homoeostasis.

## MATERIAL AND METHODS

### Mice and colonization procedures

Age-, vendor-, and strain-matched controls were used in all the experiments. C57BL/6 and BALB/c mice were obtained from Charles River. ΔdblGATA1 (GATA) mice were bred in house. A novel strain of GF GATA mice was generated by two-cell embryo transfer, as previously described^22^. Mice were bred and maintained in flexible film isolators in McMaster’s Axenic Gnotobiotic Unit. GF status was monitored weekly by DNA immunofluorescence (SYTOX Green), as well as anaerobic and aerobic culture of cecal stool samples. Mice had unlimited access to autoclaved food and water. ASF-colonized mice were originally generated by co-housing female colonizers harbouring ASF, with GF mice. ASF-colonized mice were then bred for 3 generations in individually ventilated racks within the Axenic Gnotobiotic Unit ^23^. Pathogen contamination and microbiota diversification were evaluated in mouse fecal contents every 2 weeks in sentinels by PCR for *Helicobacter bilis, H. ganmani, H. hepaticus, H. mastomyrinus, H. rodentium, Helicobacter spp*., *H. typhlonius*, and *Pneumocystis murina*. Mouse serum was also tested for murine viral pathogens by multiplexed fluorometric immunoassay/enzyme-linked immunosorbent assay (ELISA)/indirect fluorescent antibody tests ^23^. SPF colonization was performed by co-habitation of GF mice with SPF mice for a minimum of 1 month. In some experiments, mice were fed an elemental diet based on amino acids (TD1084 and TD 130916; Harlan Laboratories Inc.) for 3 generations prior to use. All procedures were approved by the McMaster University Animal Research Ethics Board.

### Intestinal cell isolation

As previously described ^24,25^, after flushing intestinal contents with cold PBS, fat was removed, and intestines were opened longitudinally and cut into approximately 3-5 mm pieces. Mucus was eliminated by washing with phosphate-buffered saline (PBS) containing 10 mM HEPES and 4 µM dithiothreitol (DTT) (Sigma) for 15 min at 37°C while on a shaker. Epithelial cells were removed by 3 rounds of 10 min-washes at 37°C, under shaking, in PBS containing 10% fetal bovine serum (FBS), 10 mM HEPES and 5 mM ethylenediaminetetraacetic acid (EDTA). The tissues were then digested in 0.125 U/mL Collagenase A (Roche) with 130 U/mL DNase I (Roche) in 10% FBS containing RPMI for 50–60 min in a shaker at 37°C. Lastly, the digested tissues were pressed through a 40 µm nylon strainer (Falcon) and immune cells were purified via 40/70% Percoll (GE Healthcare) gradient and centrifugation.

### Tissue processing and cell isolation

Bone marrow ^6^, spleen ^26^, uterus ^27^, vaginal tract ^27^, lung^28^ and blood ^6^ were collected and processed as previously described.

### EPO assay

Intestinal tissue was made up to 100 mg/mL, w/v suspension, in PBS containing complete protease inhibitors (Roche) and rotor-stator homogenized (Polytron; Kinematica, Lucerne, Switzerland). The intestinal homogenate was centrifuged (1952 g x 10 min, 4°C) and the pellet was resuspended in 2 mL of 0.2 % NaCl for 30 sec followed by 2 mL of 1.6% NaCl before centrifugation and resuspension at 100 mg/mL in Hank’s balanced salt solution (HBSS) containing 0.5% hexadecyltrimethylammonium bromide (HTAB) (Sigma: H5882). Pellets were homogenized and freeze/thawed 3 times using liquid nitrogen. Finally, samples were centrifugated, the supernatants were transferred to clean tubes and EPO activity was measured as previously described ^29^.

### Cytokine and chemokine array

Intestinal homogenates were prepared as described^30^ with minor modifications: small intestinal samples were homogenized with 20 mL/g of buffer (T-Per Tissue Protein Extraction Reagent, Thermo Scientific) containing protease inhibitors (Sigma). Samples were then centrifuged for 30 min and protein concentration was measured using the DC Protein Assay according to standardized protocols (Bio-Rad). Protein concentration in supernatants was normalized to 1 mg/mL and frozen at −80°C until assay. Cytokine and chemokine levels were determined via a cytokine 44-plex discovery assay (MD44) and a high-sensitivity 18-plex discovery assay (MDHSTC18) performed by Eve Technologies (Calgary, AB). To visualize broad differences in the metabolite signals, raw values were converted to a log scale. The fold change in protein levels between GF and SPF mice was represented on a heatmap with R software using the heatmap package. Data analysis of cycle threshold values was conducted using the Relative Expression Software Tool-384 (REST-384) version 1.

### Flow cytometry

Antibodies were obtained from eBioscience, BD Biosciences, or BioLegend. In all assays, cells were incubated with anti-FcγRII/III before incubation with fluorochrome-conjugated antibodies. Dead cells were excluded by propidium iodide uptake (Sigma) or fixable viability dye eFluor780 (eBioscience) and gated on singlets. On average, a minimum of 300,000 live and singlet cells were analyzed. Fluorescence minus one (FMO) and isotype controls were used for gating. Data were acquired on an LSR II or Fortessa (BD) and analyzed using FlowJo (Treestar). Flow-sorting experiments were performed on a FACS ARIA III (BD).

### Food allergy model

Peanut butter (3.75 mg; ∼1 mg of protein; Kraft, Northfield, IL) with 10 µg of cholera toxin (List Biologicals, Campbell, CA) in 0.5 mL of PBS was administered intragastrically (Delvo SA, Biel, Switzerland) weekly for 4 weeks. Serum was collected by retro-orbital bleeding and analyzed for peanut-specific Igs via sandwich ELISA ^31-33^.

### Histology

Intestinal segments were collected and fixed in 10% formalin for 24 h, and then washed with 70% ethanol and paraffin embedded. Sections were stained with the Protocol Hema 3 stain set (Fisher Scientific, Hampton, NH) and Masson’s Trichome method ^34,35^. For image analysis of histological sections, Masson’s trichrome stained tissue slides were scanned (VS120-ASW v2.9 slide scanner, with UPlanSApo 20x objective, Olympus) and analyzed using HALO® Image Analysis Platform (v2.2.1870.34, Indica Labs Inc, Corrales, New Mexico) using Area Quantification module (v1.0).

### Transmission electron microscopy

Immediately after excision, tissues were immersed in fixative consisting of 3% formaldehyde and 1% glutaraldehyde in 0.1-M phosphate buffer (pH 7.2). After the initial fixation, samples were post-fixed in 1% osmium tetroxide for 1 h, dehydrated in graded acetone solutions, and embedded in Polybed 812 (Polysciences, Inc.). Ultrathin sections (60–80 nm) were cut on an LKB MK III ultratome and routinely contrasted with uranyl acetate and lead citrate. The sections were examined using a FEI Tecnai Spirit BioTWIN transmission electron microscope (Fei)^6^. Eosinophil circularity was calculated as 4Π(total cell area/squared cell membrane perimeter); a value of 1.0 indicates a perfect circle. The granule content was calculated as (cytoplasm area = total cell area - nuclear area) - (the sum of all granule areas).

### Statistics

Data were analyzed and graphed with GraphPad Prism 8 software (GraphPad Software). Continuous data are expressed as means ± SEMs and were analyzed by using 1-way ANOVA with Bonferroni *post hoc* tests and unpaired Student’s *t* test. Differences were considered statistically significant at a *P* value of less than 0.05 or as indicated.

## RESULTS

### The microbiota regulates the frequency of intestinal eosinophils

In order to quantify eosinophil frequency, LP cells from the SI were isolated and evaluated by flow cytometry ^6^. Consistent with previous reports ^6,36^, intestinal eosinophils were identified as CD45^+^ SSC^high^ Siglec-F^+^ cells and expressed high levels of CD11b (**Fig. 1A**). This putative population of eosinophils was flow-sorted and stained using Hema 3 to validate their identity. The microscopic analysis showed hallmark morphological features (*i.e.* lobular polymorphic nucleus and eosinophilic granular cytoplasm) of eosinophils ^1,2^ in >95% of the cells (**Fig. 1B**).

**Figure 1.**
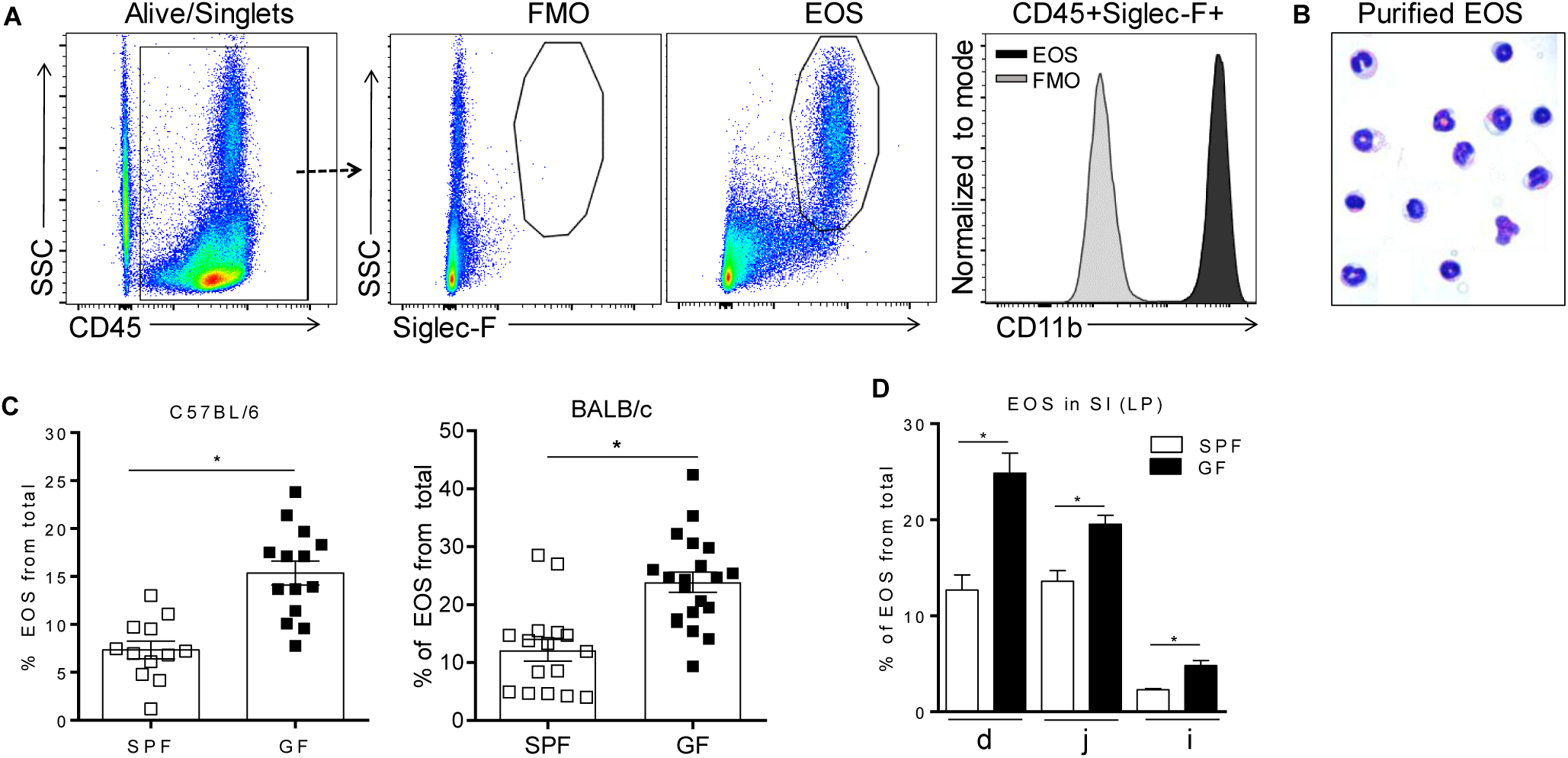
Flow cytometric identification of small intestinal eosinophils (EOS) as live singlet CD45+Siglec-F+ cells (A), morphologic validation (B) and assessment of their frequency in the small intestine (SI) of SPF and GF C57BL/6 and BALB/c mice (C, D). The frequency of EOS from total cells in the lamina propria (LP) of different sections of the SI including duodenum (d), jejunum (j) and ileum (i) of C57BL/6 mice (D). Pooled data from 3-4 experiments (n=12-20) (C) or representative data from 3 experiments (D) represented as mean ± SEM, *P<0.05

Next, we compared eosinophil frequency in the SI of SPF and GF mice. It is known that in GF mice, compared to SPF, the total mass of the SI and the total surface area are decreased, the LP is thinner and less cellular and the cell renewal rate is lower^18,37-43^. Therefore, we considered that the most informative analysis would be to focus on the frequency of immune cells. To account for possible differences in strains biased towards Th1 and Th2 immunity ^44^, both C57BL/6 and BALB/c strains were assessed. The frequency of intestinal eosinophils from total cells in GF mice was ∼2-fold higher than in SPF controls, regardless of the strain (**Fig. 1C**). We then assessed the distribution of eosinophils along the SI tract of GF and SPF mice. Regardless of colonization status, eosinophils were enriched predominantly in the proximal end of the SI (duodenum) and reduced in the distal end (ileum) (**Fig. 1D**). Nevertheless, within all sections of the SI, GF mice harbored a greater proportion of eosinophils than SPF control mice. These data demonstrate that the microbiota significantly influences the basal tissue eosinophilia of the SI LP.

### The microbiota sets the basal eosinophilic tone in naturally colonized mucosal surfaces

To test whether the differences in eosinophils between GF and SPF mice were directly related to the microbiota, we colonized GF mice with either a complex (SPF) or simple (ASF) microflora ^23^. To this end, we co-housed separate groups of GF mice with either ASF or SPF mice. The data show that the eosinophilia seen in the SI of GF mice was partially attenuated by colonization with a minimal assortment of only 8 well-defined bacterial species ^23^ using ASF mice. Eosinophilia was fully attenuated with complex colonization using SPF flora (**Fig. 2A**). These data show a graded regulatory relationship between the complexity of the microbiota and tissue eosinophil levels.

**Figure 2.**
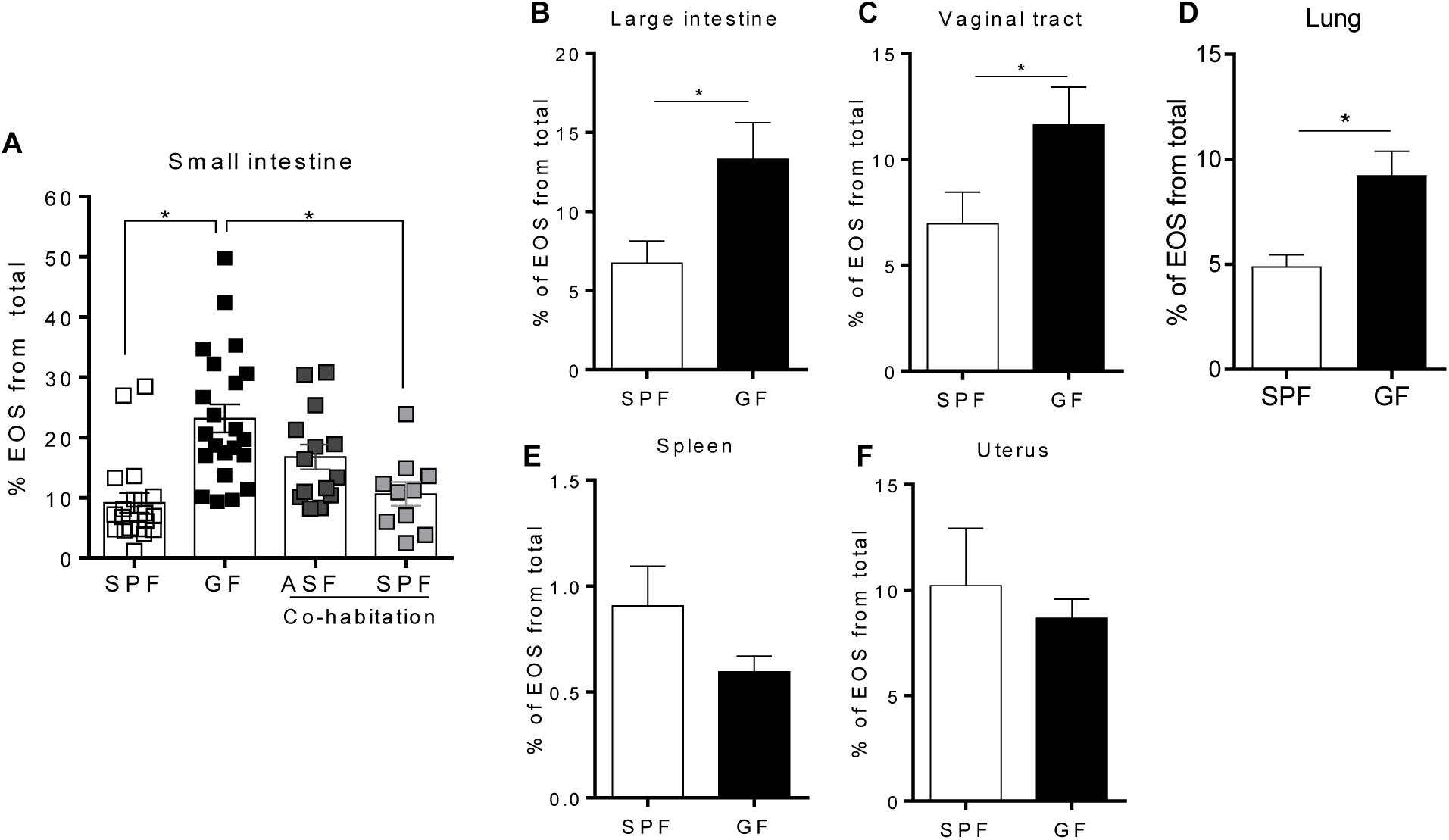
Separate groups of GF mice were colonized by co-habitation with ASF or SPF mice and the presence of small intestinal eosinophils (EOS) was assessed (A). Assessment of EOS frequency in the large intestine, vaginal tract, lung, spleen and uterus of SPF and GF mice by flow cytometry (B-F). Pooled data from 2-5 experiments (n=8-22) represented as mean ± SEM, *P<0.05.

Since the microbiota regulated the frequency of enteric eosinophils, we next examined whether GF induced eosinophilia was present in other naturally colonized mucosal sites (large intestine, vaginal tract and lung), sterile -or poorly colonized-mucosal sites (uterus) ^45^ and non-mucosal sites (spleen). The frequency of eosinophils was also significantly higher in the large intestine, vaginal tract and lung of GF mice compared to SPF (**Fig. 2B-D**) but not in sterile tissues such as the spleen (**Fig. 2E**) and uterus (**Fig. 2F**). These data further support the concept that the frequency of tissue eosinophils is dependent on the natural colonization status.

### The maturity of the immune system is not associated with intestinal eosinophilia

It is well established that the adaptive immune system of GF mice is immature, particularly as it refers to T cells ^46^. Consistent with previous observations ^47,48^, we found that the frequency of CD4^+^ T cells was significantly lower in the SI of GF mice compared to SPF controls (**Fig. 3A**). In contrast, there were no statistically significant differences in the frequency of B cells, dendritic cells (DCs), macrophages and mast cells (**Fig. 3A**). To investigate whether the eosinophilia observed in GF conditions was due to the absence of colonization or inherent to an immature immune system, we generated microbiota-competent mice with an immature immune system ^49^. To this end, BALB/c mice, fed with an elemental (amino acid) diet and housed in a SPF environment, were bred for 3 generations. Fewer Peyer’s patches and a lower frequency of CD4^+^ T cells were observed in the LP, thus confirming the immaturity of the adaptive immune system of these mice (**Fig. 3B**). We did not find changes in the frequency of B cells, macrophages or eosinophils. These findings indicate that the relative eosinophilia in GF mice is likely due to a lack of microbial-derived signals rather than immune immaturity *per se*.

**Figure 3.**
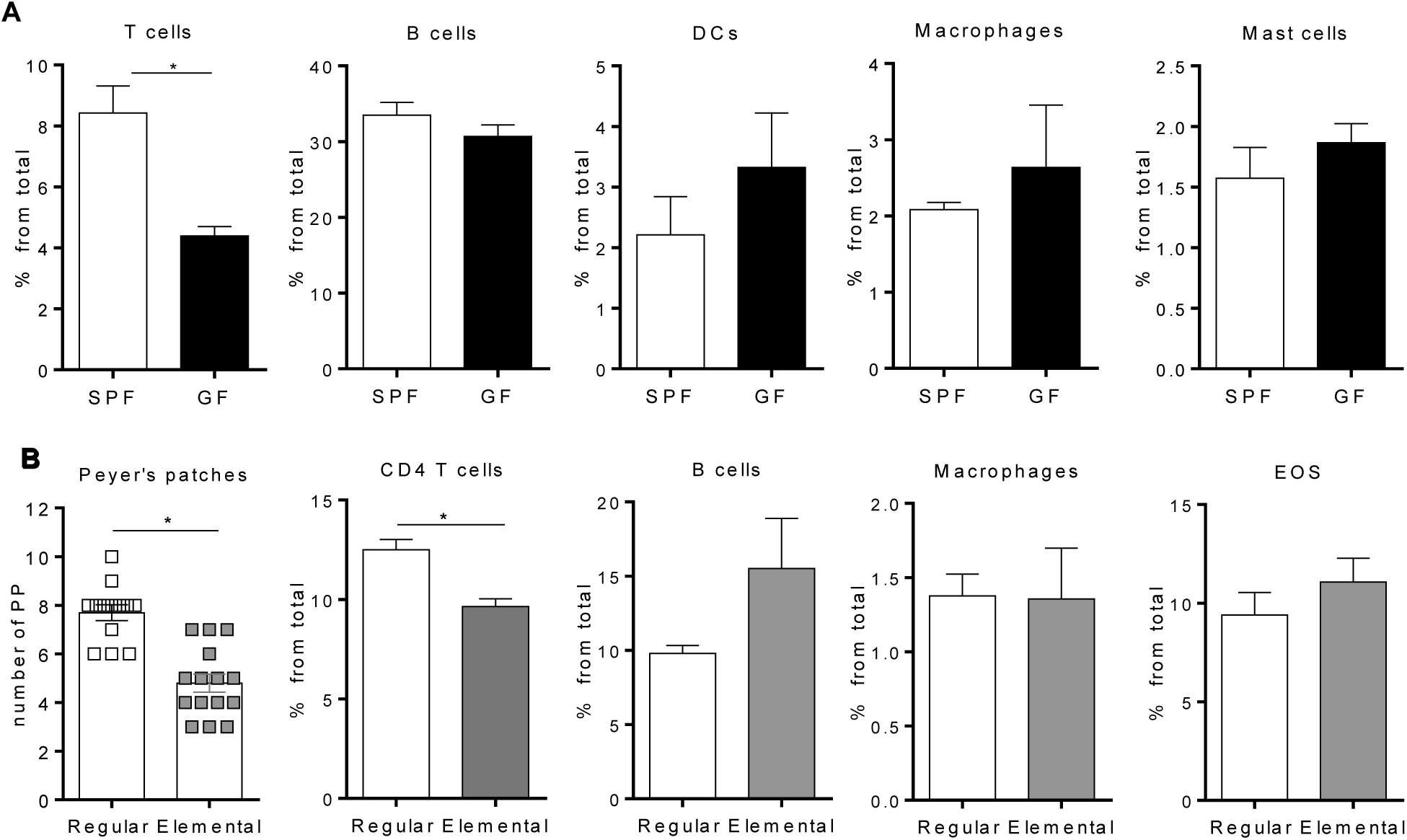
Flow cytometric characterization of adaptive and innate immune compartments (A, B) in the small intestine of GF and SPF mice fed a regular (A) or elemental (B) diet (B). Pooled data from 2-4 experiments (n=6-22) represented as mean ± SEM, *P<0.05.

### The microbiota regulates eosinophil attraction and retention signals in mucosal tissues

Tissue accumulation of eosinophils associated with chronic intestinal inflammation is typically attributed to increased eosinopoiesis^50^. Here, we investigated whether the same mechanism applies to constitutive eosinophilia in healthy GF mice. We found that there were no differences in BM eosinophils between GF and SPF mice, nor were differences in EOS circulating in the peripheral blood (data not shown). This suggested that eosinophil accumulation in mucosal sites of otherwise healthy GF mice might be mediated by increased expression of attraction and/or retention signals in the SI, large intestine, lung and vaginal tract.

To identify signals associated with eosinophil migration and retention at the tissue level, we analyzed chemokine and cytokine levels in the proximal (duodenum) and distal (ileum) SI segments of GF and SPF mice (**Fig. 4A**). Overall, cytokine and chemokine production was lower in the SI of GF mice as compared to SPF; this is consistent with the underdeveloped mucosal immune system reported in GF mice^18^. Nevertheless, compared to SPF controls, GF mice exhibited a significant increase in IL-3^51^ and VEGF^52,53^, which are associated with eosinophil chemotaxis and survival **(Fig. 4B**). Furthermore, we observed a significant decrease in the levels of IL-11^54^ and CXCL9^55^ in GF mice, which regulate eosinophil recruitment (**Fig. 4B**). These data suggest that the microbiota regulates signals involved in the attraction, retention and survival of eosinophils in the SI, the absence of which results in the relative eosinophilia found in GF conditions.^51^

**Figure 4.**
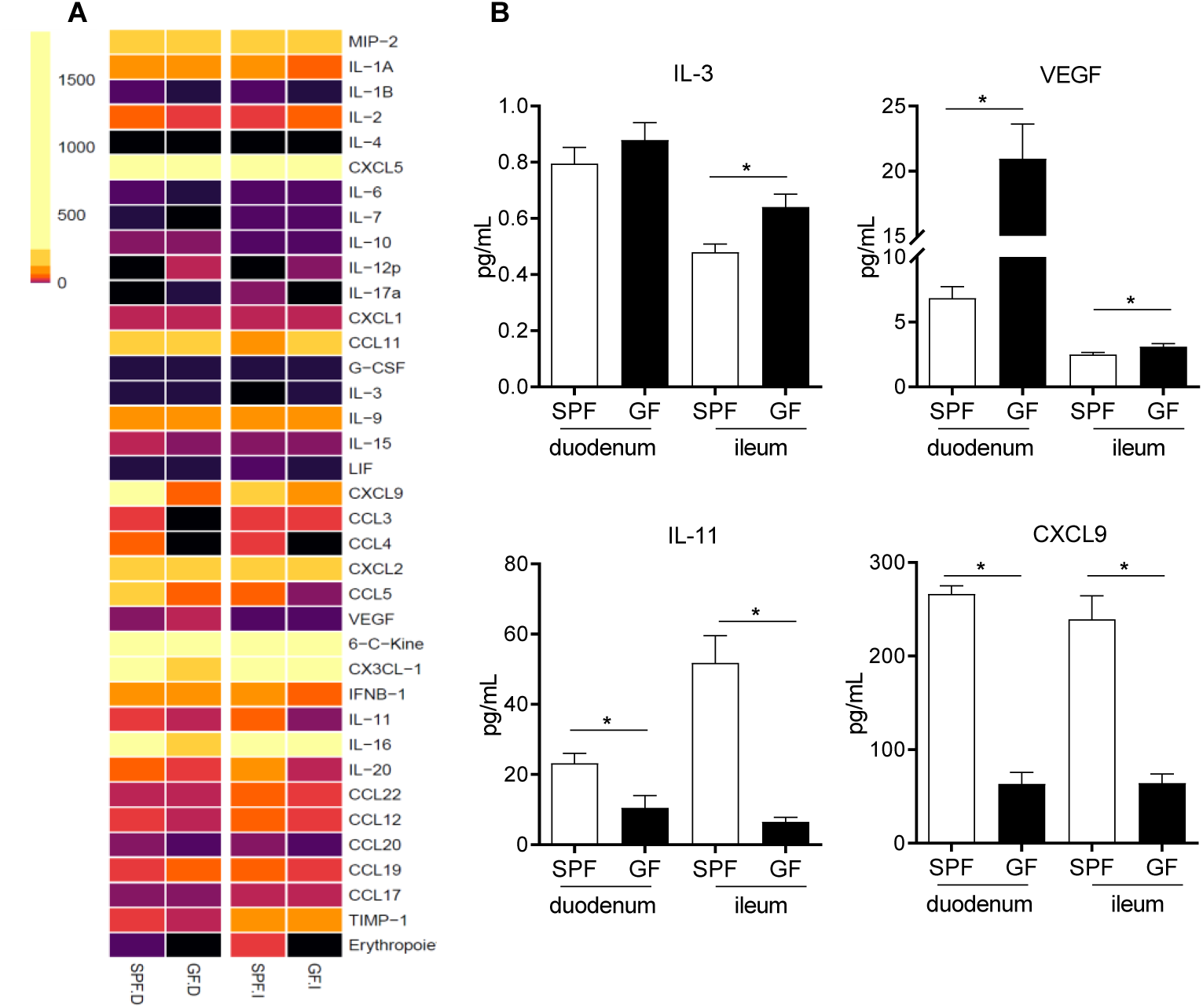
Heatmap of chemokine and cytokine protein levels in proximal (duodenum; D) and distal (ileum; I) small intestinal homogenates from GF and SPF mice as determined by protein array (A). Relevant proteins differentially produced both in the proximal and distal small intestine of SPF and GF mice (B). Data from 6 mice (A, B) represented as mean ± SEM, *P<0.05.

### Eosinophil morphology and functionality are influenced by the microbiota

In order to evaluate eosinophil function in GF mice, we performed comprehensive transmission electron microscopy analysis of small intestinal sections from GF and SPF mice (**Fig. 5A-D**). Eosinophils in GF mice (**Fig. 5B-D**) exhibited profound morphological alterations in comparison to SPF controls (**Fig. A**). Eosinophils from GF mice had significantly more cytoplasmic protrusions; this was quantified and measured as reduced circularity of the outer membrane perimeter (**Fig. 5E**). Signs of eosinophil cytoplasmic lysis (ECL), potentially associated with the cytoplasmic protrusions, were found, although very rarely, in eosinophils from GF mice (**Fig. 5D**). While increased cytoplasmic protrusions are indicative of cell activation, we did not find evidence of eosinophil degranulation in either of them. Strikingly, intestinal eosinophils from GF mice had significantly less granule content (**Fig. 5F**) and granules of smaller size than SPF counterparts. To substantiate these findings, the degranulation status of eosinophils was also determined by quantifying EPO activity ^29,56^ in SI homogenates of GF and SPF mice. Despite the extensive eosinophilia (**Fig. 1C**), significantly lower EPO activity was detected in GF mice (**Fig. 5G**) compared to SPF controls.

**Figure 5.**
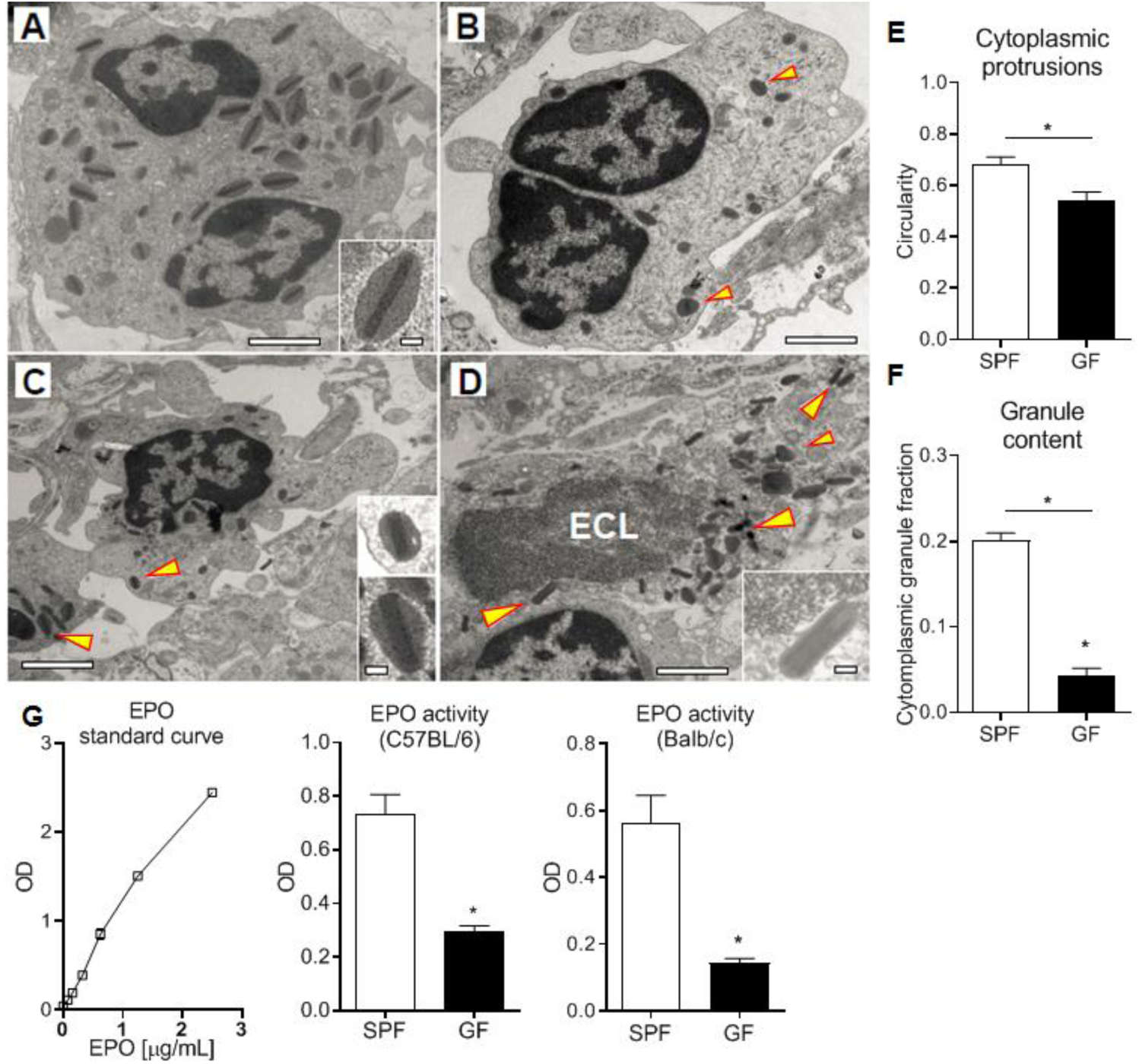
Normal SI eosinophils transmission electron microscopy ultrastructure (A), showing bi-lobed nuclei and a high density of granules composed of an electron-dense core surrounded by an electron-lucent matrix rich in EPO (arrowhead). SI eosinophils from GF mice (B-D) exhibit cytoplasmic protrusions, measured as reduced circularity (E), lower content of cytoplasmic granules (F) and, occasionally, ECL signs (D). Assessment of EPO in intestinal homogenates of SPF and GF mice (G). Representative (A-D) and pooled (E-F) data from 6 mice, and pooled data from 2 independent experiments (n=6) (G) represented as mean ± SEM, *P<0.05.

**Figure 6.**
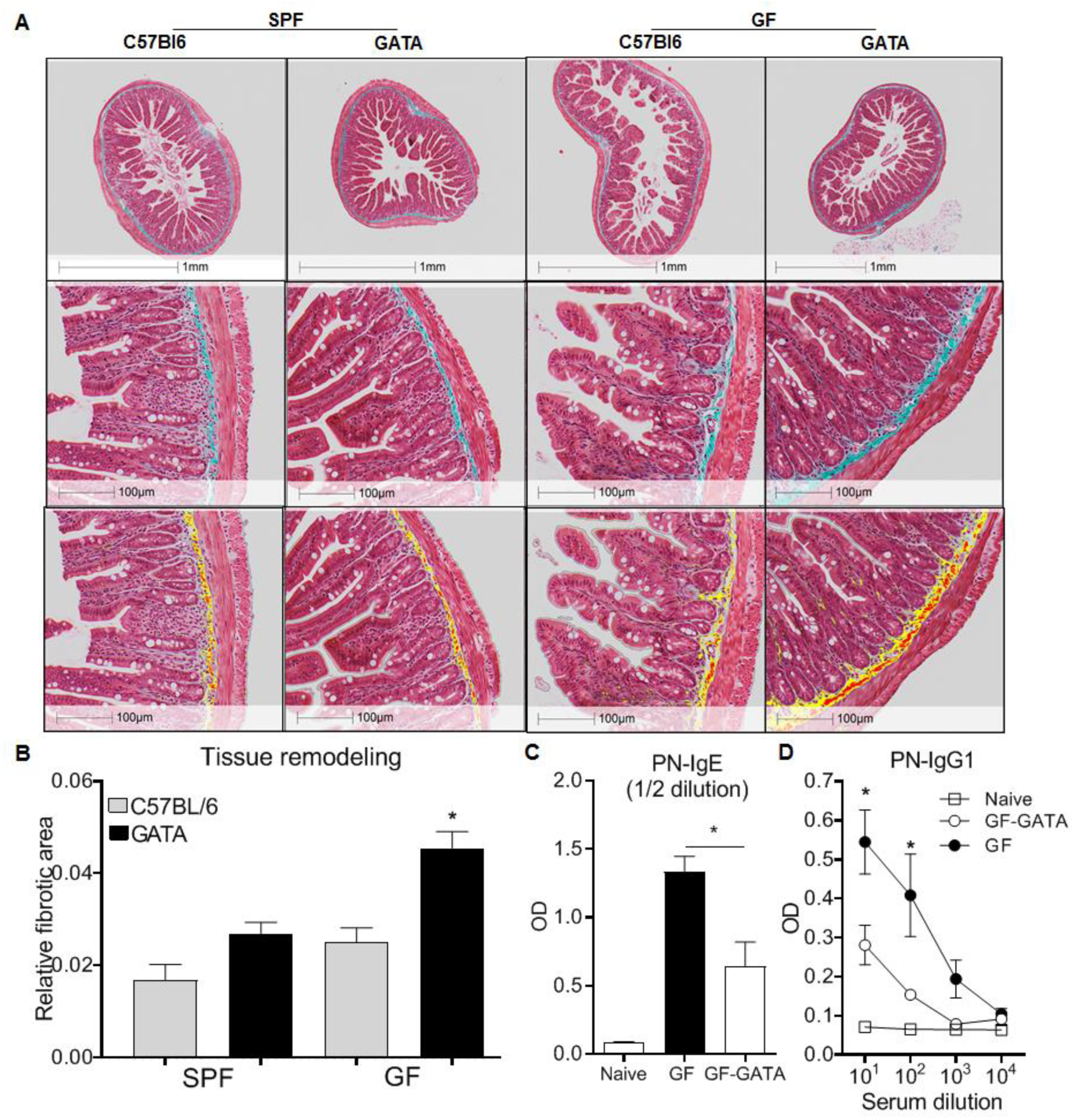
Representative histological examples of intestinal fibrosis (A, upper panels) and associated Halo quantification of fibrotic surface on digitalized (20X) sections where yellow and red represents Masson’s Trichrome positive stain (A, lower panels) and associated quantification (B) of overall fibrotic area performed on GF mice, eosinophil deficient GF mice (GATA-GF) and controls. GF mice, eosinophil deficient GF mice (GATA-GF) and controls were intragastrically sensitized to peanut (PN) and serum levels of PN-specific IgE (C) and IgG1 (D) were determined by ELISA. Data from 4-5 mice (A, B) or pooled data from 2 experiments (n=6) (C, D) represented as mean ± SEM, *P<0.05.

### The microbiota modulates eosinophil-mediated intestinal remodeling and allergic sensitization

To investigate the functional significance of the morphological features observed in GF mice, we generated a novel eosinophil-deficient mouse strain (ΔdblGATA; GF-GATA) on a GF background. Given that eosinophils are known to contribute to tissue remodeling, intestinal tissues were assessed with Masson’s Trichome staining ^34,35^ to evaluate collagen deposition (**Fig. 5A**). To remove potential bias of field selection, these differences were quantified using the HALO Image Analysis software. This analysis demonstrated higher collagen density in the submucosal layers of eosinophil-deficient mice, particularly in GF conditions, as compared to SPF controls **(Fig. 5B).** These findings suggest that the absence of eosinophils in GF GATA mice is associated with increased fibrotic tissue remodeling.

Intestinal eosinophils play a critical role in the initiation of allergic responses to food allergens. We have previously demonstrated that eosinophil-deficient (ΔdblGATA) mice are unable to produce peanut-specific immunoglobulins^6^. Additionally, gut dysbiosis has been associated with a higher risk of developing allergy in humans ^57-60^, and GF mice have been shown to be prone to generate Th2 responses ^61,62^. In this context, we evaluated if the lack of microbial modulation of eosinophils may impact allergic sensitization to foods. We employed an established food allergy model using intragastric sensitization to peanut^24,32,63,64^ in GF mice on a GATA or C57BL/6 backgrounds. In agreement with previous studies^62^, GF mice generated higher levels of peanut-specific IgE and IgG1 than SPF controls (**Fig. S1**). However, the production of both immunoglobulins was significantly lower in GF mice deficient in eosinophils as compared to GF controls **(Fig. 5C-D)**. These findings raise the possibility that the microbiota influences the development of food allergic sensitization partly through its effects on intestinal eosinophils.

## DISCUSSION

Mucosal eosinophils have been traditionally considered recruited proinflammatory cells, whose biological benefit is limited to defence against parasitic infections. However, new evidence has established that eosinophils also contribute to the initiation, propagation, and resolution of innate and adaptive immune responses, and to homeostatic tissue repair and remodeling ^4,5,65,66^. The diversity of functions ascribed to enteric eosinophils in particular has propelled investigation into the mechanisms that influence their tissue residence and functionality. In this regard, the microbiota is known to be critical for the development of the SI mucosal adaptive immune system ^18^. Given that eosinophils natively inhabit the SI ^67^, we investigated the impact of the microbiota on intestinal eosinophil frequency and function.

Comprehensive quantification of intestinal eosinophils was carried out by flow cytometry ^6,36^ and further validated by morphological analysis (**Fig. 1A, B**). The total frequency of intestinal eosinophils in either C57BL6 or BALB/c was ∼2 fold higher in GF mice as compared to SPF controls (**Fig. 1C**). This is in contrast with the findings by Mishra *et al.* ^68^, who did not observe significant differences in eosinophils numbers in the GI of SPF and GF mice when quantified by histology. Several factors might have contributed to this discrepancy including the strain of mice (Black Swiss mice *vs* C57BL/6 and BALB/c), microbiome differences under SPF conditions, the number of mice utilized in each study (n=5 *vs* n=12-20) and, lastly, the technique employed for eosinophils quantification, as it is likely that flow cytometry allowed for a more comprehensive and precise quantification than immunohistochemistry using the eosinophil granule protein, major basic protein (MBP) ^69^. Importantly, we show that intestinal eosinophils in GF mice contain have granules of smaller size and less cytoplasmic granule content than SPF controls, which may lead to eosinophil underdetection when using MBP-based methods.

Consistent with previous reports ^6,68^, eosinophils were pre-eminently localized in the proximal end of the SI and decreased in frequency distally. However, the proportion of eosinophils in each section was significantly higher in GF mice compared to SPF controls (**Fig. 1D**). Importantly, colonization of GF mice with a complex microbiota reduced enteric eosinophils to a frequency comparable to that in SPF mice. This shows that enteric eosinophilia is greatly influenced by the host intestinal microbiome (**Fig. 1E**).

We next considered whether enteric eosinophilia could be an innate response to compensate for the immaturity of the adaptive immune system that occurs in GF mice. It is known that feeding SPF mice with an elemental diet results in an immature immune system ^70^. Our data show that the administration of an elemental diet for 3 generations resulted in features indicative of immune immaturity, such as a reduced number of Peyer’s patches and lower CD4^+^ T lymphocytes. However, it did not lead to the eosinophilia identified in GF mice fed with a conventional diet (**Fig. 2**), which suggests that enteric eosinophilia is independent of the maturity of the adaptive immune system and thus dependent on the microbiota.

The relative eosinophilia observed in GF mice, its regulation by complex microbial colonization and its independence of the maturity of the adaptive immune system, along with the findings of unperturbed eosinopoiesis suggest that the microbiota directly regulates enteric eosinophils through interactions with cells resident in the mucosal compartment. The significant changes in the production of signals (IL-3^51^, VEGF^52,53^, IL-11^54^ and CXCL9^55^) that regulate eosinophil migration, attraction and survival of eosinophils in the SI of GF mice supports this notion (**Fig. 4B**).

The formation of cytoplasmic protrusions might relate with a migratory stage and is indicative of cell activation; despite their abundance in eosinophils from GF mice (**Fig. 5E**), there were no signs of degranulation. In fact, EPO levels were lower in the SI of GF mice as compared to SPF (**Fig. 5G**), consistent with the smaller content and size of cytoplasmic granules shown by transmission electron microscopy (**Fig. 5F**). The drastic morphological changes observed in eosinophils from GF mice, as compared to SPF, evidence that aspects related to granule processing and maturation, as well as functionality, are influenced by the microbiota. While eosinophil activation is often associated with tissue damage, eosinophils also participate in tissue homeostasis^72^. For example, increased eosinophil activation has been detected in patients with ulcerative colitis (UC), as compared to controls. Interestingly, the number of activated eosinophils was shown to be greater during the remission phase of UC ^73^, which may suggest a dual role in intestinal inflammation and repair. We found that in mice lacking both eosinophils and microbiota, collagen deposition in the submucosal layer was double to that of mice deficient in either eosinophils or microbiota alone (**Fig. 5B, C**), suggesting that at least certain components of tissue remodeling are regulated by interactions between the microbiota and eosinophils.

Several lines of evidence have indicated that dysbiosis in humans is associated with an increased prevalence of allergic sensitization ^57,59,74-76^. However, the mechanisms underlying this association remain to be fully elucidated. It was recently shown that GF and antibiotic-treated mice developed increased allergic sensitization as compared to SPF mice ^62^. On the other hand, we have shown that enteric eosinophils are essential to the induction of allergic sensitization in SPF mice^6^. These observations raise the question of whether the microbiota and enteric eosinophils synergize in the induction of allergic sensitization. Here, having demonstrated that GF mice exhibit heightened eosinophilia, we show that the absence of eosinophils in GF mice results in an attenuation of allergic sensitization. Thus, these data support the concept that the microbiota influences the capacity to develop allergic sensitization, at least in part, through its effects on enteric eosinophils.

In summary, this study demonstrates that eosinophil frequency and activation in the intestinal mucosa is regulated by the microbiota. It also shows that processes such as tissue repair and the induction of allergic sensitization appear to be regulated by an interplay between the commensal microbiota and intestinal eosinophils. Given that the tissue microenvironment crucially shapes the nature and evolution of subsequent antigen-host interactions, these data have fundamental implications to understanding the role of the microbiota and eosinophils in health and disease.

## AUTHOR CONTRIBUTIONS

RJS conceptualized the project and designed experiments. RJS and VA performed experiments, analyzed the data and wrote the manuscript. TW, MEG, TSM, RC, JFK, YE and AH helped with experiments. HJG conducted experiments in the Axenic and Gnotobiotic Unit. EFV generated GATA-GF mice. SA and KA performed HALO analysis. JE conducted EM analysis. EFV, KA and DKC provided scientific input and revised the manuscript. MJ obtained funding, oversaw the project and edited the manuscript.

## FUNDING

This work was supported by Food Allergy Canada, the Delaney family, the Zych family, the Walter and Maria Schroeder Foundation and AllerGen NCE. EFV holds a Canada Research Chair and is funded by CIHR grant MOP# 142773.

## ACKNOWLEDGMENTS

We acknowledge Dr. Puja Bagri for critical review of the manuscript. We thank all the members of the Jordana Lab for technical help and scientific input.

